# Multi-block RNN Autoencoders Enable Broadband ECoG Signal Reconstruction

**DOI:** 10.1101/2022.09.07.507004

**Authors:** Michael Nolan, Bijan Pesaran, Eli Shlizerman, Amy Orsborn

**Affiliations:** Department of Electrical and Computer Engineering, University of Washington, Seattle WA 98195; Center for Neural Science, New York University; Department of Applied Mathematics, University of Washington; Department of Bioengineering, University of Washington

## Abstract

**Objective:** Neural dynamical models reconstruct neural data using dynamical systems. These models enable direct reconstruction and estimation of neural time-series data as well as estimation of neural latent states. Nonlinear neural dynamical models using recurrent neural networks in an encoder-decoder architecture have recently enabled accurate single-trial reconstructions of neural activity for neuronal spiking data. While these models have been applied to neural field potential data, they have only so far been applied to signal feature reconstruction (e.g. frequency band power), and have not yet produced direct reconstructions of broadband time-series data preserving signal phase and temporal resolution.

**Approach:** Here we present two encoder-decoder model architectures - the RNN autoencoder (RAE) and multi-block RAE (MRAE) for direct time-series reconstruction of broadband neural data. We trained and tested models on multi-channel micro-Electricorticography (*μ*ECoG) recordings from non-human primate motor corticies during unconstrained behavior.

**Main Results:** We show that RAE reconstructs micro-electrocorticography recordings, but has reconstruction accuracy that is band-limited to model scale. The MRAE architecture overcomes these time-bandwidth restrictions, yielding broadband (0-100 Hz), accurate reconstructions of *μ*ECoG data.

**Significance:** RAE and MRAE reconstruct broadband *μ*ECoG data through multiblock dynamical modeling. The MRAE overcomes time-bandwitdh restrictions to provide improved accuracy for long time duration signals. The reconstruction capabilities provided by these models for broadband neural signals like *μ*ECoG may enable the development of improved tools and analysis for basic scientific research and applications like brain-computer interfaces.

## 1. Introduction

Measurements of brain activity collected for neuroscience experiments and neural engineering applications, such as electrocorticographs (ECoG), local field potentials, multiunit spiking activity, are complex, multidimensional time series. Recordings from a given modality exhibit nonlinear dynamics [1] and non-stationary signal statistics [2]. Each neural data modality has different statistical properties [3], which must be accounted for in neural analyses and neural information decoding methods.

Modeling methods provide a means to simplify neural recording complexity. In particular, neural signal models assume *a priori* structure for recordings that allow identification of signal and noise. Dynamical models further aim to relate sequential time series data across different temporal scales. Classical modeling methods either linearly relate features of neural activity to sensorimotor stimuli or neural recording values; or build these relations through a dynamical system with *a priori* defined models [4, 5, 6, 7, 8, 9]. Linear methods benefit from preconditioning steps such as feature estimation and trial response averaging that reduce the variance in neural recordings and find task- or stimulus-specific average responses. These analyses regard inter-trial variability as noise, discarding useful information from individual neural recordings. More recent methods avoid trial averaging and feature engineering by fitting models to learned distributions of single trial neural data [10, 11]. Recurrent neural network (RNN) autoencoders such as LFADS model neural recordings as an observed projection from a latent dynamical system [12]. LFADS produces spike rate estimates that reliably decode task parameters without the need for task parameter averaging or *a priori* signal feature computations. Analysis at the single-trial level has provided new insights into neural processing, and improved our ability to decode behavior from neural activity [11].

Since different neural measurement modalities produce signals with varied statistics, methods to build dynamical models for a wide range of signals are needed. Encoder-decoder networks like LFADs have primarily been applied to spiking data, but ECoG and other field potential recordings notably differ from the multiunit spiking (MUA) recordings. Spiking data is a discontinuous point process, whereas field potentials are a continuous varying signal. Field potentials like ECoG, in particular, are potentially challenging to model because they are spatially and temporally rich. ECoG contains broadband information spanning a wide range of frequencies at varying power scales, and represents larger spatial scales than spiking activity while also capturing highly local spiking activity [13, 3, 14]. These properties also make ECoG a promising modality for neural engineering applications [15, 16].

RNN autoencoders have been applied to model continuous neural data including ECoG band power [17], but have not been used to directly model the underlying time-series. Broadband ECoG signal power features have been accurately modeled by low-dimensional dynamical factors with trajectories effectively separating distinct behavioral states [17]. ECoG band power features and other continuous data that have been modeled with LFADS like calcium imaging fluorescence [18] and rectified electromyography (EMG) [19] are considerably less variable than the ECoG time-series. Pre-specified feature estimates from rich data like ECoG may discard information that could be more fully leveraged by directly modeling the time-series itself. For instance, ECoG power features exclude phase that may contain important information [20]. Enhanced models that can accurately reconstruct ECoG signals may ultimately lead to improved technologies for decoding information from neural data.

RNN autoencoder models like LFADS—a subset of more general encoder-decoder RNN latent modeling methods—have not been used to directly model ECoG time-series data. We set out to use encoder-decoder models to reconstruct a dataset of *μ*ECoG signals recorded from multiple motor cortical areas of a freely behaving primate over a four day period. While ECoG data is difficult to directly reconstruct with a dynamical model because of its nonstationary variability and dynamics, RNN encoder-decoder models provide a degree of expressiveness that may enable direct ECoG signal reconstruction. The reconstruction task serves as an optimization objective that causes the model to learn the dynamics in the dataset, and also serves as the basis for neural signal processing applications like data completion and anomaly detection that will advance neurotechnologies.

Here we demonstrate two new encoder-decoder models and apply them to the ECoG reconstruction task. Our first model, the RNN Autoencoder (RAE), was able to partially reconstruct individual ECoG trials but failed to accurately reconstruct high frequency ECoG signal components (<60Hz). We found that reconstruction bandwidth, the range of constituent signal frequencies accurately estimated in a reconstruction task, decreased as trial length increased. Our second model, the Multiblock RNN Autoencoder (MRAE), overcame these time-bandwidth limitations. MRAE separates the input signal into different frequency bands and separately reconstructs each filtered signal component. The encoded activity is then mixed between model blocks to create an estimate of the full input signal. We found that MRAE more accurately reconstructed direct ECoG signals than RAE for long reconstructions while retaining a high reconstruction bandwidth. MRAE’s broadband ECoG reconstruction capabilities will enable improved single-trial and real-time analysis of ECoG signals and other rich neural time-series data.

## 2. Methods

### 2.1. Data Collection and Preprocessing

All analyses were performed on neural activity recordings collected from motor cortical areas of a rhesus macaque (*macaca mulatta*, male, 9.5 kg, 8 years old) over a period of four days (Figure 1). All procedures were approved by the Animal Care and Use Committee at New York University. Recordings spanned free-behavior periods of 22 hours each day. During these periods, the subject was allowed to behave without instruction, reward, presented stimulus or penalty within the confines of their enclosure. Data were collected using Blackrock Wireless neural recording platform [21, 22]. We used this dataset when training and evaluating our models due to their long recording duration, which provided us with a large training data volume.

**Figure 1.**
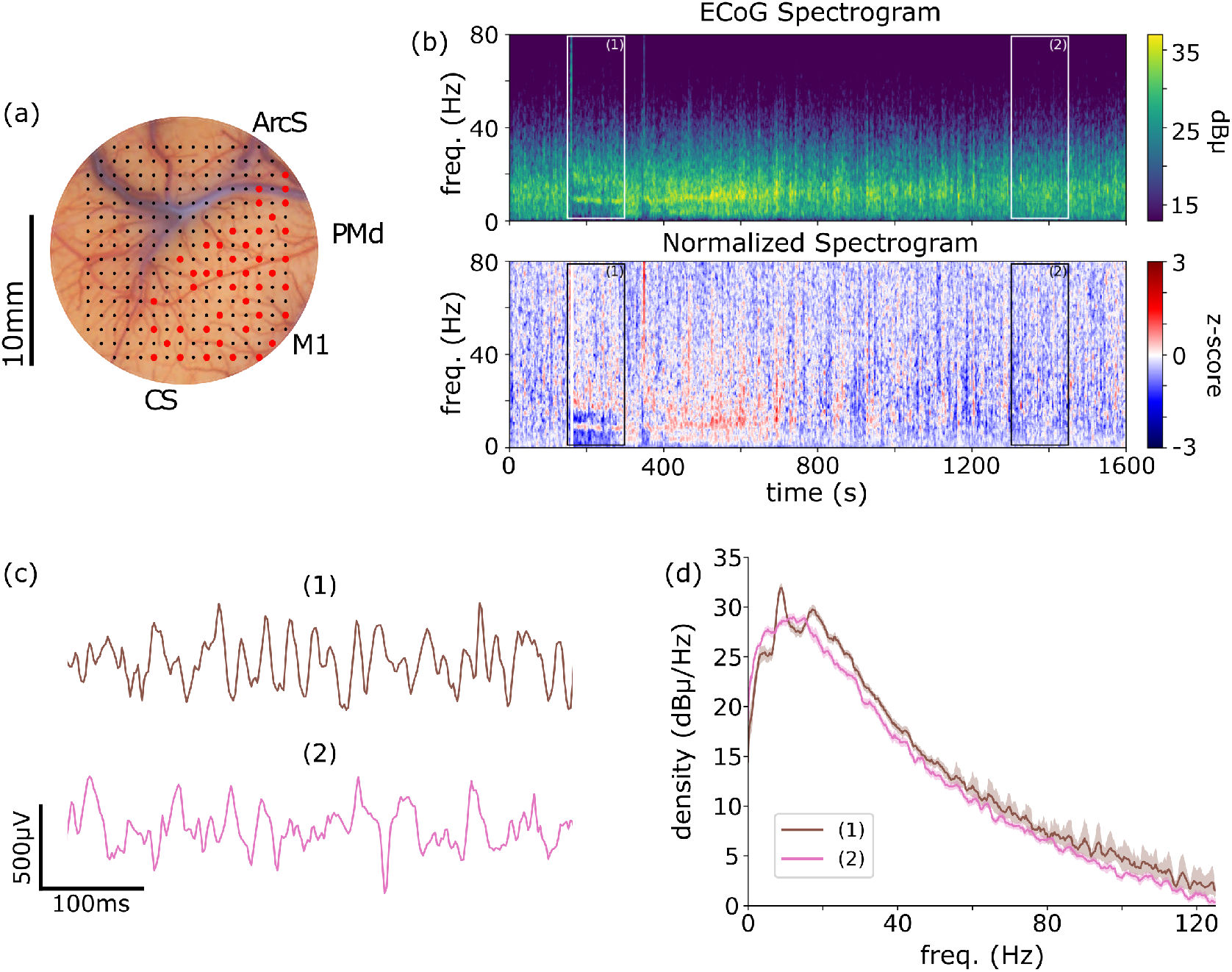
ECoG dataset profile. (a) ECoG electrode placement. Full array of *μ*ECoG electrodes are shown overlayed on the subject’s cortical surface. Electrodes used in this report’s analyses are labeled in red. M1: Primary motor area. Abbreviations: PMd: Dorsal premotor area. ArcS: Arcuate sulcus. CS: Central sulcus. (b) Spectrogram and normalized spectrogram. Normalized spectrogram is z-scored within each computed frequency bin. (c) Traces from highlighted spectrogram regions. (d) PSD from highlighted spectrogram regions.

Data were initially collected at 30kHz on 62 channels covering dorsal pre-motor (PMd) and primary motor (M1) corticies (Figure 1(a)). The dataset was then down-sampled to 250Hz and filtered to remove signal artefacts. Data channels were filtered by mean signal power to remove dead channels, rejecting data from those with power below 50% of the median power across all channels in each 2 hour segment of the dataset. The resulting dataset contained *μ*ECoG signal data from 41 of the original 62 channels and included a cumulative 33 hours of *μ*ECoG data after filtration and data rejection. The channels used in this analysis are indicated by red markers in Figure 1. When reconstructed, data were segmented into non-overlapping trial windows. We normalized each data segment (trial) by z-scoring to a mean of zero and a standard deviation of 1 for each channel.

### 2.2. Reconstruction Task and Performance Metrics

The reconstruction task requires the dynamical model to estimate an input target sequence *x* (”trg”) with an output estimate sequence 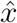 (”est”). The task objective is to minimize the reconstruction error, which is measured as the mean square error (MSE) between the target and the estimate:

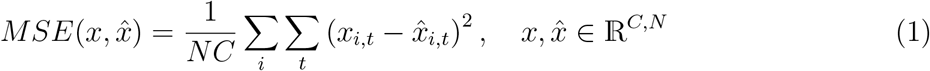

where *N* is the number of time points in each sequence and *C* is the number of ECoG channels. Index variables *i*, *t* iterate over channel and time values, respectively. Models are optimized to minimize the MSE of their reconstructions using the adaptive momentum (ADAM) algorithm. [23]. Default optimization hyperparameters were used from the pytorch implementation [24]. MSE is also used as a reconstruction performance metric. A reconstruction error of 0 is ideal, while a reconstruction error of 1 is of equal magnitude to the variance in the signal segment due to data normalization.

#### 2.2.1 Reconstruction Bandwidth

In addition to MSE, reconstruction accuracy was assessed in terms of signal frequency bandwidth. The bandwidth of a reconstruction was determined to be the range of frequency space in which the model output signal was sufficiently similar to the target signal. We estimated signal reconstruction bandwidth (BW) by computing the PSD of the error signal 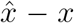 and counting the number of quantized frequency bins over which the error was sufficiently low power (less than −3dB, i.e. half the power of the target signal). This count was then scaled by the frequency bin width to create a bandwidth estimate. All PSD estimates were computed using a single-trial multitaper PSD with a fixed frequency bin width of 5Hz [25]. Normalized PSD estimates or reconstruction residues (difference between reconstruction and target) were referenced to the mean PSD of the target data sequences. Negative normalized PSD values indicate accurate reconstructions while 0dB output values indicate that the reconstruction residue was equal to the target signal at that frequency and therefore indicates an inaccurate fit at that frequency band.

### 2.3. Models: RAE and Multiblock RAE (MRAE)

We developed two encoder-decoder generative models to reconstruct motor cortical *μ*ECoG signals. These models are similar in type to Latent Factor Analysis for Dynamical Systems (LFADS) models proposed for spiking data [12, 11]. While the type of the models is similar, in order to address *μ*ECoG signals, the models that we introduce differ by including adaptations to apply them to *μ*ECoG signals and by introducing additional components for precise reconstruction as described below. The first model, RAE, directly models neural field potential signals as diagrammed in Figure 2. This model is modified from the original LFADS architecture in a manner reflecting the differences between spiking and neural field potential data. The output of RAE model directly estimates the *μ*ECoG signal at each time point, while the original LFADS model estimated the poisson process firing rate most likely to generate the observed spiking data in a given trial. In RAE, the original LFADS Poisson negative log-likelihood optimization objective is replaced with a mean-square error (MSE) objective computed between the model output and the input ECoG signal. [17].

**Figure 2.**
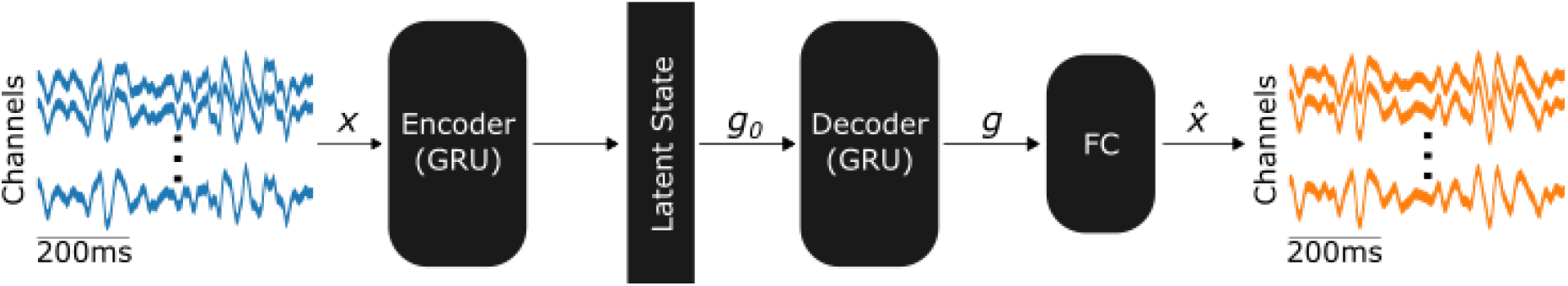
Single-block RAE model architecture. The bidirectional GRU encoder block estimates latent variable distribution parameters from the entire *μ*ECoG input trial. A latent state value is sampled from this distribution and then used to initialize the hidden state of a unidirectional encoder GRU. This decoder output is then passed through a linear output layer to create an output signal estimate. Network parameters are optimized by minimizing MSE loss between the estimate and the input signal.

Analysis of RAE reconstruction performance indicated that direct adaptation was insufficient to model broadband neural field potential recordings. To address this shortcoming, we propose a novel multiblock, parallelized extension of RAE called Multiblock RAE (MRAE, diagrammed in Figure 3). MRAE is a parallelized form of the RAE model architecture which spectrally decomposes the input into non-overlapping frequency bands. The bands of filtered inputs are then individually reconstructed with a separate RAE block. The broadband signal is fit by training the model to mix the concatenated generator outputs of all model blocks into a single multichannel signal estimate through a fully-connected network layer. Each block is optimized according to its own individual reconstruction loss, while the generator output mixing layer is optimized to minimize the broadband reconstruction error. These learning objectives are in MSE estimates, as is the case of the single-block RAE discussed previously.

**Figure 3.**
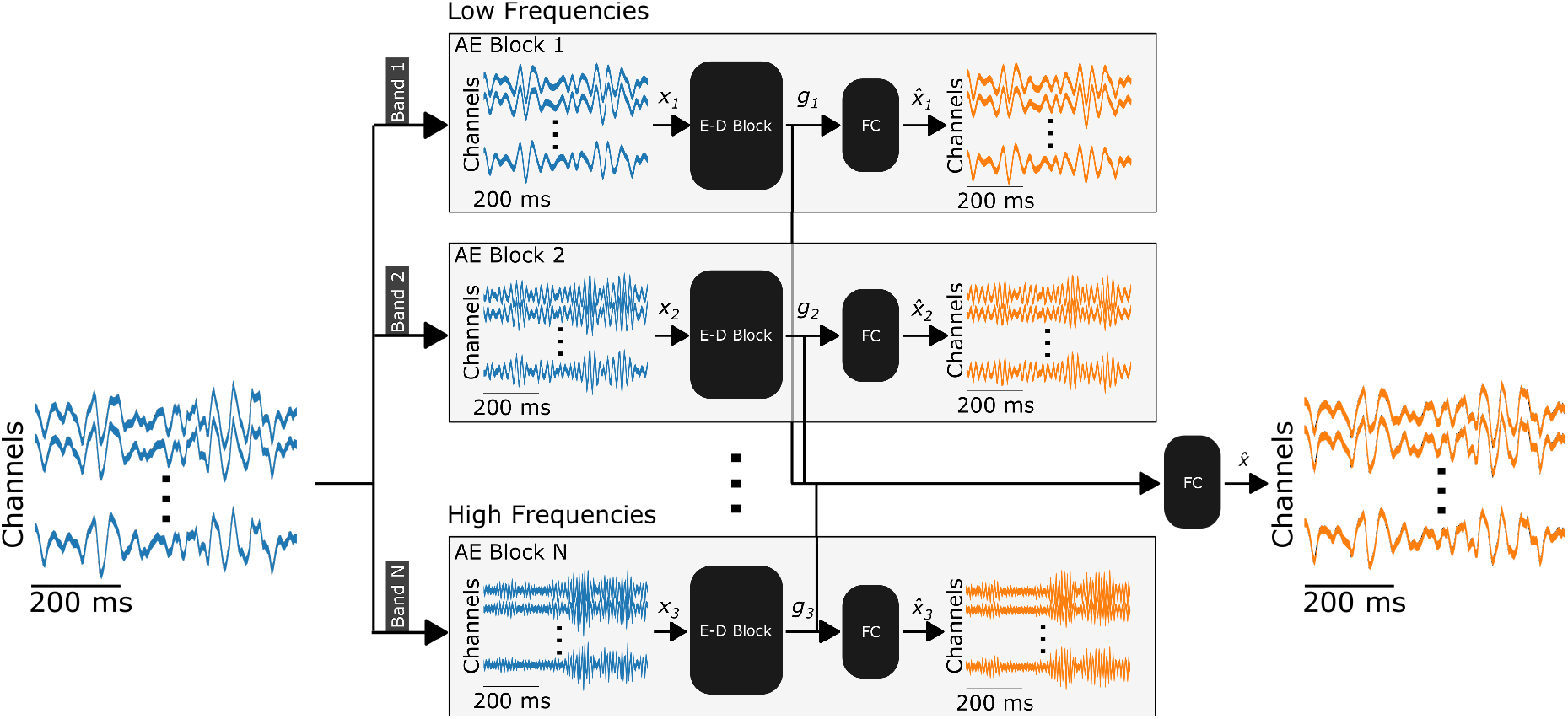
Multiblock RAE (MRAE) model architecture. Input trials are bandpass filtered into non-overlapping bands that uniformly span the full bandwidth of the input. Filtered inputs are then reconstructed by independent autoencoder blocks. Decoder outputs *g*_*i*_ are concatenated before projection through a last linear layer (FC) to estimate the broadband input target. Individual blocks are optimized independently to minimize bandlimited signal reconstruction error. FC is optimized to minimize broadband reconstruction error.

## 3. Results

### 3.1. ECoG Reconstruction with RAE and MRAE

ECoG signal reconstruction results from both the RAE and MRAE model architectures are shown for a single 600ms trial in Figure 4. Reconstructions are shown for the mean channel which is computed by averaging the signal across all channels for each time point independently. This example shows that RAE model outputs were accurate, but band-limited. This limitation of the RAE model was observed consistently across all trials. The RAE model accurately reconstructed low frequency signal components, but was unable to accurately reconstruct high frequency signal components. MRAE *μ*ECoG reconstruction, shown in Figure 4-bottom, retained low-frequency reconstruction performance while also more accurately reconstructing high-frequency signals.

**Figure 4.**
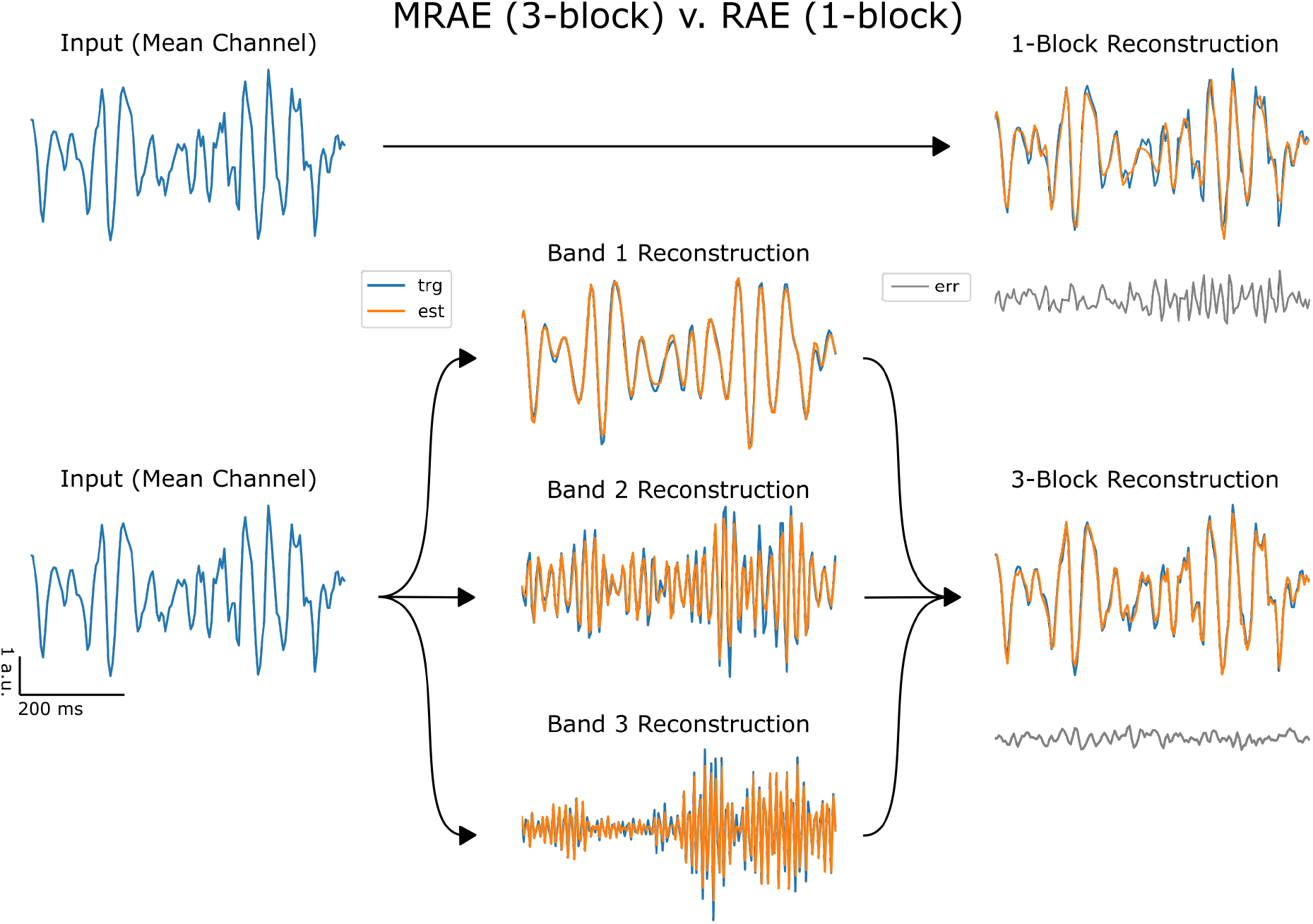
ECoG signal reconstruction example. A single 600ms trial is shown with the target signal (trg) in blue and the reconstruction estimate (est) in orange. Reconstruction error traces (err) are plotted in grey. Mean channel reconstructions are shown for the single-block RAE model (top) and 3-block MRAE model (bottom). Individual block reconstructions are shown in the bottom center sequence of axes. Blocks are arranged vertically and descend in accordance with increasing frequency bands.

### 3.2. Spectral Characteristics of ECoG Reconstructions

Our findings further indicated that signal reconstruction bandwidth varies with the length of the reconstruction task. This is shown in Figure 5(a)). Comparing mean power spectral density (PSD) of target signals for both 1-block and 3-block model reconstructions revealed differences in the reconstruction bandwidth. More closely overlapping PSD plots indicate that the reconstruction has a similar bandwidth as the target signal. We found that the 1-block model reconstruction bandwidth decreased as the length of the reconstruction increased. In contrast, the 3-block model reconstruction bandwidth covered the target signal bandwidth for all task sequence lengths up until 1000ms length (Figure 5(a), bottom). For sequence lengths longer than 1000ms, the reconstruction bandwidth began to break down even for the 3-block model. Each individual block, trained to fit its given frequency band, only reconstructed from the lower edge of that band to an upper limit frequency less than the upper limit of the entire band. This leads to the segmented PSDs observed for the 1000ms trial.

**Figure 5.**
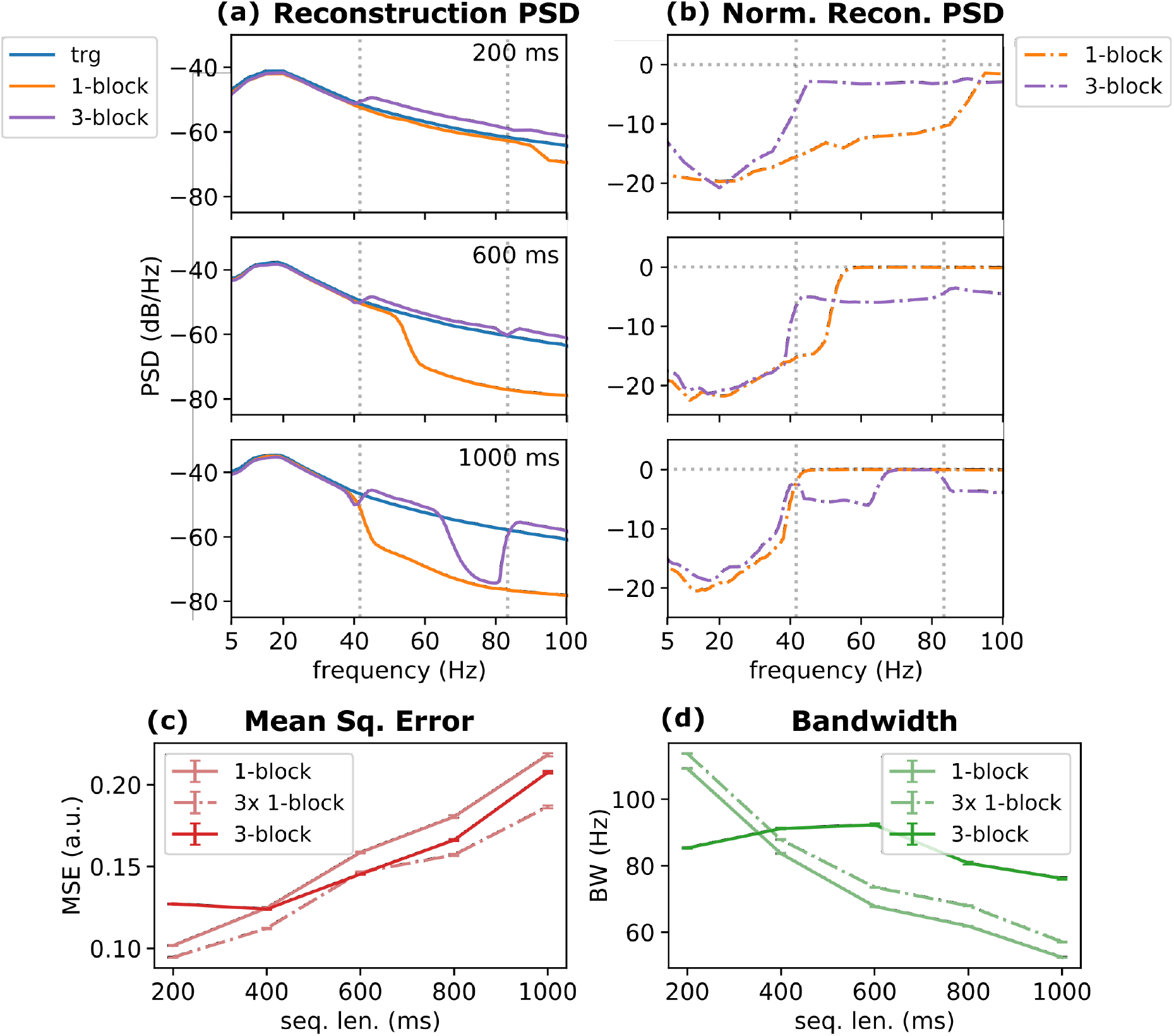
Mean reconstruction power spectral density (PSD) and reconstruction performance for 512-unit RAE and 3-block MRAE models. (a) Mean target PSD plots of ECoG mean-channel signals are shown with reconstruction PSDs. 1-block (orange) and 3-block (purple) models are shown. Frequency ranges are limited to their low frequency resolution limit and high frequency limit defined by the decimation filter used during downsampling to 250Hz (100Hz). (b) Reconstruction error PSDs, normalized by the target signal PSD, are shown for 1- and 3-block models. 0 dB/Hz normalized error corresponds to a residual signal having equal power density to the target signal at that frequency. Vertical dotted lines denote multiblock frequency band edges. Horizontal dotted lines in normalized error PSD plots denote a 0dB reference. (c) Reconstruction MSE for RAE and 3-block MRAE models across all trial lengths. (d) Reconstruction bandwidth for RAE and 3-block MRAE models across all trial lengths. In (c, d), a 1-block model with 1536 units is trained for all task lengths and is plotted with the 512-unit RAE and 3-block MRAE model results.

These results reveal the band-limited nature of model reconstructions from a single autoencoder block and motivates decomposing the signal into multiple bands. We further quantified this by studying the mean PSD estimate of the residual error signal (Figure 5(b)). The normalized reconstructed error PSDs show the same pattern of increased reconstruction error and decreased bandwidth as in Figure 5(a). In particular, reconstruction bandwidth decreased as the target signal length increased. This is clear in the 1-block RAE model (orange) across all signal lengths. The MRAE model reliably fit the lowest frequency band (i.e. inputs to block 1) accurately. Higher frequency bands, reconstructed from the outputs of higher MRAE blocks, produced a higher normalized error, though these reconstructions had a lower error than the reconstructions produced from the 1-block model. The combined effect of processing each band individually in the MRAE model was to reduce the overall reconstruction error at longer reconstruction lengths when compared to the 1-block RAE model.

### 3.3. Reconstruction Performance Across Target Sequence Lengths

To quantify the supported bandwidth of the models that we consider, we compared reconstruction performance in the time-domain using MSE and reconstruction bandwidth (BW) (see Methods) for the 3-block MRAE and 1-block RAE models across reconstruction task trial lengths (5(c)). MRAE had a lower reconstruction for all target reconstruction lengths of 400ms or greater. RAE outperformed MRA on average on trials with 200ms signal lengths. A similar pattern of model performance was observed for reconstruction bandwidth (5(d)). While 1-block reconstruction bandwidth decreased with increased trial target length, 3-block MRAE reconstructions maintained a high reconstruction bandwidth. These results affirm the trend of error bandwidth increasing with trial length shown in the right column of Figure 5(b).

One possible explanation for the improved performance of MRAE over RAE could be a difference in model sizes, since by adding blocks, the MRAE increases the total unit count relative to the RAE. To exclude the explanation that the better performance of MRAE is merely due to its larger size, we fit a 1,536-unit (3 x 512) RAE model to each reconstruction task length that matched the total unit count used in the MRAE model tested (Figure 5(c, d), dotted lines). Increasing the unit count in the RAE models did lead to improved MSE performance, both relative to the MRAE and smaller RAE. However, increasing the unit count did not eliminate the reconstruction bandwidth limitations, which are observed in both the 512-unit and 1,536-unit RAE models. Performance data is also summarized in Tables 1 and 2 for MSE and BW, respectively.

**Table 1.**
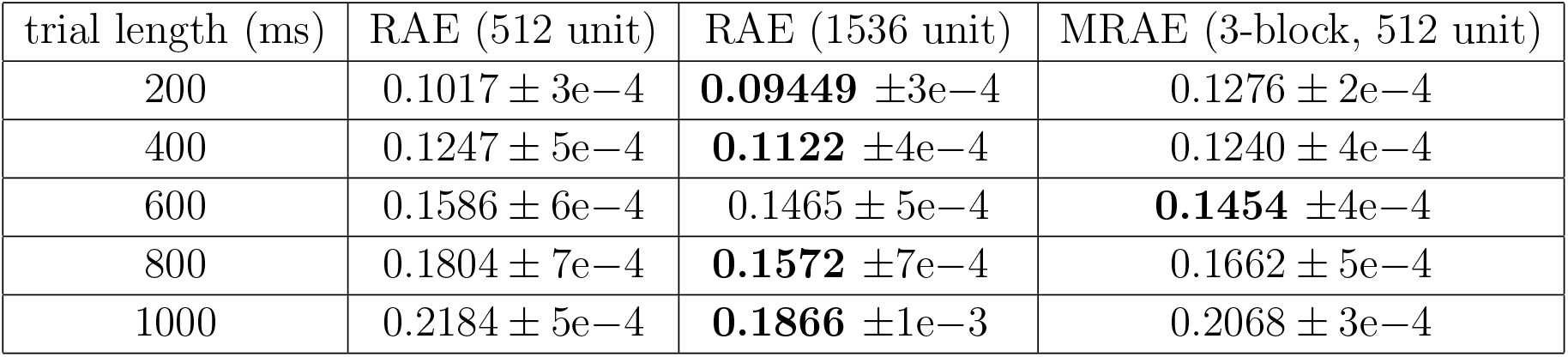
Mean squared error (MSE) ablation table across RAE, 3x unit count RAE and 3-block MRAE. Mean ± 95% CI half-width. Best performance bolded for each trial length.

**Table 2.** Bandwidth (BW) ablation table across RAE, 3x unit count RAE and 3-block MRAE. Mean ± 95% CI half-width. Best performance bolded for each trial length. High reconstruction bandwidth indicates an accurate reconstruction, with a maximal value of 125Hz imposed by the 250Hz sampling rate.

| trial length (ms) | RAE (512 unit) | RAE (1536 unit) | MRAE (3-block, 512 unit) |
| --- | --- | --- | --- |
| 200 | $109.3 \pm 3e-1$ | **$113.8 \pm 6e-2$** | $85.34 \pm 2e-1$ |
| 400 | $83.65 \pm 1e-1$ | $87.82 \pm 1e-1$ | **$91.19 \pm 2e-2$** |
| 600 | $67.73 \pm 1e-1$ | $73.51 \pm 1e-1$ | **$92.30 \pm 3e-1$** |
| 800 | $61.82 \pm 1e-1$ | $67.91 \pm 1e-1$ | **$80.72 \pm 4e-1$** |
| 1000 | $52.40 \pm 1e-1$ | $57.09 \pm 1e-1$ | **$76.17 \pm 3e-1$** |

## 4. Discussion

In this work we presented novel neural data RNN autoencoder models, 1-block RAE and 3-block MRAE, and demonstrated their application to multiple ECoG signal reconstruction tasks. RAE is modeling ECoG recordings as a dynamical system that reconstructs signal activity, however is limited in supporting wide bandwidths when reconstructing longer sequences. To overcome time-bandwidth limitations in this model, we introduce a multiblock recurrent autoencoder, MRAE. We show that MRAE sustains its reconstruction bandwidth over a larger range of reconstruction task lengths than RAE and we also show that MRAE reconstructs ECoG data more accurately than RAE.

While we find that the multi-block architecture can improve reconstruction performance, there are some exceptions to this pattern. Notably, the RAE model outperforms the 3-block MRAE model when reconstructing very short (200ms) ECoG trials (Figure 5(c, d)), with both better reconstruction error and bandwidth. However, as trial length increased, the accurate frequency range of the RAE outputs quickly narrows. MRAE, in contrast, produces residual errors with more consistent bandwidth as trial length increases. However, these are slightly lower accuracy than those of the RAE for short reconstruction lengths.

Our results highlight that each RAE block has a fixed reconstruction performance ceiling (Figure 5). For a fixed model size, increasing trial length resulted in increased reconstruction error and decreased reconstruction bandwidth. This pattern indicates that a single RAE block will have a fixed time-bandwidth output capacity that serves as an upper limit to model performance in time series reconstruction tasks. The MRAE model overcomes this limit by distributing the signal bandwidth over multiple blocks. For all trial lengths shorter than 1000ms, individual block reconstruction capacity was not exceeded by the input signal bandwidth and reconstructions were accurate across the block’s entire frequency band (Figure 5). At 1000ms trial lengths, individual block reconstructions failed to span their entire frequency band, leading to frequency components at the upper end of each band not being reconstructed by model outputs. Our results highlight limitations of RAE methods that must be considered carefully when analyzing neural data. Models like LFADS have primarily been applied to model features that vary on relatively slow timescales, such as multi-unit firing rates [11], calcium fluorescence imaging [18], field potential power features[17], and rectified electromyography signals [19]. Our results demonstrate the challenges of extending existing methods to data with richer dynamics, and provides a new architecture for modeling broadband neural data.

Importantly, the block architecture of MRAE provides an efficient way to improve performance and badnwidth in time-series reconstruction. We found that increasing RNN size improves model reconstruction accuracy (MSE), but that bandwidth limitations are better addressed by our multi-block architecture. For RNN models, parameter counts scale as the square of the total unit count [12]. Adding units to an RAE improves performance, but at a high cost of memory required for model implementation. In contrast, the multiblock model (MRAE) total parameter count scales linearly with the number of blocks used. This enables accurate reconstruction over a wider frequency band with an overall smaller model than an similarly performing single block RAE. This provides some important context to the performance patterns seen in Figure 5(c, d) and Table 1, where the RAE model has better reconstruction accuracy performance than the MRAE when RNN unit counts are equal. This performance boost comes with a 10x increase in the total number of parameters. This accuracy gain does not accompany a similar gain in reconstruction bandwidth, where MRAE consistently outperformed RAE. The improved scaling of the MRAE architecture’s reconstruction bandwidth with reconstruction lenth may be particularly important when considering applications like closed-loop neural interfaces that require low-latency, real-time processing of longer continuous *μ*ECoG sequences. Similar systems requiring direct dynamical modeling of shorter *μ*ECoG sequences would benefit from the single-block RAE model, which did show better reconstruction accuracy and bandwidth at the shortest tested sequence length (200ms).

A more complete understanding of the source and the nature of time-bandwidth capacities will be critical for improving the generality of autoencoder modeling methods for neuroscience. Dynamical modeling efforts toward other neural recording modalities (e.g. LFP, calcium imaging) must account for their broadband spectral distributions. The time-bandwidth limit we observed shows that both RAE and MRAE models are biased to reconstruct low-frequency signal components more accurately than high-frequency signal components. While MRAE enables a wider range of reconstructed frequencies than RAE, each model block will reconstruct the lower end of its input frequency band more accurately than frequencies at that band’s higher end. The causes for this limitation may come from multiple sources. The inductive bias of GRU RNNs may be partially responsible. Other sources may be the power spectral density characteristics of ECoG and the optimization objective used to train the model. The power spectra of ECoG and other neural field potential signals follow a 1/*f* distribution. When the model is optimized using the L2 norm as an error objective, accounting for larger amplitude deviations in low frequencies are prioritized by the model. During model development, we explored using alternative training objective functions such as L1 norm and a log-scale frequency domain error that might be less biased towards low frequencies. However, we found that the L2 norm consistently produced models with the most accurate signal reconstructions out of all objective functions tested. While additional exploration is needed, we hypothesize the superior performance of L2 norm training objectives may be attributable to the nonconvexity of the L1 norm and frequency-domain error metrics. Future exploration of alternate training objectives and model architectures will be important for further improving autoencoder methods in neuroscience.

The RAE and MRAE models we introduce here are influenced by the LFADS variational RNN autoencoder design [12, 11, 18, 19]. These types of models are capable of inferring low-dimensional dynamical latent factors that accurately estimate firing rates from multi-unit firing rate data and are useful for behavioral decoding [26, 27]. We extend this approach to develop models that can accurately reconstruct ECoG time-domain signals of variable length, segmented arbitrarily from a dataset of task-free primate motor cortical data. LFADs and MRAE models are both variants of a larger set of sequence-to-sequence autoencoder RNN models (seq2seq) that are trained to reconstruct time-series data, and learn low-rank subspaces in the process. Autoencoder networks optimized for time-series reconstruction have also been shown to *organize* latent spaces, for instance forming clusters relevant to data context, which enables unsupervised learning of data structure. Autoencoder models like MRAE developed here that can capture rich broadband data will therefore provide a tool to learn latent embeddings and brain-behavior relationships from *μ*ECoG and other broadband neural measurements.

We anticipate that the multi-block approach may also be beneficial for reconstruction tasks of other signal modalities such as multiunit spiking activity (MUA) and other neural field potential recordings (e.g. LFP). MRAE also provides opportunities for future cross-modal neural data analysis by incorporating blocks for each modality of interest. This would enable new modeling of cross-scale interactions between local (e.g. neuronal firing activity) and more mesoscale or global(e.g. ECoG, LFP, EEG) measures of neural activity. Multi-modal extensions of MRAE may also provide new ways to perform joint modeling of neural and behavioral data [28, 29, 30]. Further extensions of the multiblock architecture may also be needed to incorporate cross-band interactions to capture within or cross-modal interactions, such as cross-frequency coupling and other known neural bispectral phenomena [31].

RAE and MRAE also fit into a larger context of dynamical systems modeling in neuroscience research and neural engineering applications. The RAE and MRAE model may be useful for applications such as filling in missing measurements (data completion) or predicting future neural activity (forecasting). Reconstruction with MRAE is also a possible means of anomaly detection, where out-of-distribution neural recordings may be detected by their poor reconstruction accuracy [32]. Accurate dynamical models of neural signals also enable analysis of neural dynamics using nonlinear systems analysis, which has shed light on many complex cognitive processes [33]. This work in this report develops a novel highly accurate reconstructive dynamical model of broadband ECoG time series data. This will enable the expansion of dynamical systems methods and analysis to direct modeling of mesoscale measures of neural activity that show great promise for brain-computer interfaces and other neural devices [15, 16].

## 5. Conclusion

Neural dynamical models enable single-trial neural data reconstructions. This capability enables the latent analysis of individual trials and the flexible estimation of neural activity without reduction into predefined features or aggregation into trial statistics. Here we present two neural dynamical models based on a variational autoencoder architecture: RAE and MRAE. We show that both models provide highly accurate reconstructions of ECoG data collected over a four day period from a freely behaving primate subject. These reconstructions are band limited in the RAE experiments, favoring low frequency reconstruction over high frequency reconstruction. We further show that MRAE overcomes this limitation by expanding reconstruction bandwidth. Both models provide accurate reconstructions over a range of trial lengths up to 1.0s. We propose that these models will enable new data analysis methods including anomaly detection and data completion.

